# Volitional Attention to Color: Breaking Willed Attention out of the Spatial Domain

**DOI:** 10.1101/2025.07.10.664245

**Authors:** John G. Nadra, Grace Sullivan, George R. Mangun

**Author notes:** Correspondence to: John Nadra, Ph.D.

## Abstract

Attention can be guided by either voluntary (top-down) or involuntary (bottom-up) influences. In real-world vision, voluntary attention can be directed by a wide range of influences such as reward, priming, meaning, experience, selection history, and volition. Laboratory studies of voluntary attention have commonly used attention-directing cues to induce or instruct human subjects to attend one class of stimuli while ignoring others. In recent years, free choice conditions of attention have been introduced, where self-generated volitional shifts of attention (willed attention) are possible (for a review, see Nadra & Mangun, 2023). Prior willed attention studies have focused on mechanisms of covert spatial attention, investigating the neurophysiological correlates of willed attention related to both post-decision processing, and the pre-decision brain activity patterns that predict participants’ free choices about where to attend. Much less work, however, has been done on willed non-spatial attention. In this study, in a fashion similar to prior willed spatial attention studies, we investigated willed attention to stimulus color. In a trial-by-trial cueing design, subjects were either cued what color to attend or were allowed to freely choose what color to attend, in order to subsequently discriminate visual features of the cued or willfully chosen target. Behavioral measures showed that the subjects were selectively attending the cued or willed color. Using support vector machine decoding on EEG alpha (8-12 Hz) signals, we found significant post-decision differences in alpha-band power and broadband EEG voltage when participants chose to attend to orange versus purple. However, in contrast to the findings in prior work on willed spatial attention, no pre-decision EEG signals could be identified that predicted which color participants would choose to attend to.

## Introduction

The deployment of attention can be driven by both top-down (voluntary) and bottom-up (involuntary) sources. Voluntary, or goal-based, attention is commonly investigated using spatial cueing paradigms, in which cues (e.g., an arrow) are used to direct participants’ attention to the location(s) of relevant target stimuli that require a decision and response (Posner, 1980, 2016). The study of willed attention builds upon these methods by including cues or *prompts* that signal the participant to freely choose where to focus their attention in a given instance (Taylor et al., 2008; Hopfinger et al., 2010; for a review, see Nadra & Mangun, 2023). Although willed attention has primarily been studied in the domain of spatial attention (Bengson et al., 2014, 2015, 2020; Gmeindl et al., 2016; Liu et al., 2017; Nadra et al., 2023, 2025; Rajan et al., 2019), there are compelling theoretical grounds to suggest that it reflects a more general attentional mechanism.

Non-spatial forms of attention contribute to our perception of the world and have been studied extensively in behavioral and cognitive neuroscience research. Behavioral work has shown that selective attention to color results in improved perception and performance of the attended color stimuli independent of spatial attention (Ansorge & Becker, 2014; Brawn & Snowden, 1999; Lee et al., 2018; Kasten & Navon, 2008; Kingstone, 1992), and that color signals in visual search arrays can be both potent cues for identifying task-relevant targets (Ansorge et al., 2010; Folk & Remington, 1998) and as powerful distractors when irrelevant (Theeuwes & Burger, 1998). Cognitive neuroscience studies have shown that attention to color modulates sensory responses to color stimuli (Andersen & Hillyard, 2024; Anllo-Vento et al., 1998; Corbetta et al., 1991; Diepen et al., 2016; Eimer, 1997; Lee et al., 2018; Noah et al., 2020). Since color is a powerful cue for human perception (Hansen & Gegenfurtner, 2009), and like spatial locations can serve to direct goal-directed attention (Theeuwes, 2025), it is reasonable to ask whether willed color attention operates by similar mechanisms as willed spatial attention.

One key finding in willed spatial attention research is that the pattern of EEG alpha signals over posterior scalp regions in the few hundred milliseconds <u>prior</u> to a free decision about where to focus covert attention is predictive of the participants’ subsequent choice (Bengson et al., 2014). A recent study by Wang and colleagues (Wang et al., 2024) investigated willed color attention in a paradigm where colored moving dots served as the target stimulus, and participants were either cued or willfully chose in advance of the appearance of the dot arrays which color was relevant on each trial. They probed EEG alpha power in the period prior to the appearance of the choose cues, but did not find any evidence of predictive patterns in the EEG, suggesting that willed spatial attention and willed color attention may depend on different neural mechanisms in the pre-decision period. In the <u>post</u>-decision period, that is after a choice cue, Wang and colleagues observed many of the same neural responses as were reported in Bengson and colleagues (2015).

There were, however, several differences in the design of the study compared to prior willed spatial attention studies that could have contributed to the negative result for pre-decision EEG alpha predictive signals. First, the target stimuli were presented for 2000 ms, or more than twenty times as long as in Bengson and colleagues (Bengson et al., 2014). In addition, the cue-to-target stimulus onset asynchrony (SOA) and the inter-trial-interval (ITI) were both much shorter (1000-2000 ms versus 2000-8000 ms) in the study by Wang and colleagues. Finally, the probability of receiving a choice trial was higher in their study (50% versus 33%). These differences may have induced the subjects to adopt strategies that differed in fundamental ways from those used by participants in prior willed spatial attention studies. For example, the long stimulus durations used by Wang and colleagues might have resulted in the subjects delaying their willed decisions until the stimulus arrays appeared, and as a result, antecedent brain states that might have influenced rapid decision choices at appearance of the choice cues might then have contributed little to biasing choices as to what color to attend. This in no way a criticism of Wang et al (2024). To further test whether willed color attention may share neural mechanisms with willed spatial attention, we conducted a study that matched many task parameters with the original 2014 study of Bengson, including stimulus duration, SOA and ITI. In addition, we developed a more challenging color discrimination task that, in principle, should have encouraged subjects to develop a strong attention set during the choice trials.

## Methods

### Stimuli and Procedure

Thirty healthy participants were recruited for this study (26 females, 4 males). One subject was removed for inability to follow task instructions, two others were removed for having target discrimination accuracy lower than the criterion of 75%, and four more were removed for excessive EEG artifacts (artifact rejection exceeding 25% of their data). Thus, twenty-three participants were retained for final analysis.

We implemented a cueing paradigm similar in many features to prior willed spatial attention designs (Figure 1). A fixation point was present at the center of the screen for the entire duration of the experiment, and participants were instructed to maintain ocular fixation on this point throughout the study. An auditory spoken cue or choose prompt of the words ‘orange,’ ‘purple,’ or ‘choose’ began each trial. For the color cues, the subjects were instructed to prepare to discriminate the cued color by focusing their attention on that color. On choose trials they were instructed to freely and spontaneously choose where to attend. Thus there are two main conditions in this paradigm, with the ‘orange’ or ‘purple’ cues denoting the ‘instructed (cued) condition,’ and the ‘choose’ prompt denoting the ‘willed (choose) condition’ where participants freely chose which of the two colors to attend. Following the cue or prompt was an SOA varying between 2-8 seconds, then a target appeared which consisted of an array of nine lines where each line was randomly tilted 45 degrees to the left or right. Participants were instructed to press a button signaling whether the line of the attended color was tilted to the left or right. A report screen was then presented for 3,000 ms with the word ‘Color?’ presented above the fixation point. This required that the subjects report whether they attended to orange or purple on that trial. The addition of the report screen allows for an analysis of the willed condition, but also acted as a validation that participants were paying attention to the cues in the instructed condition; participants correctly reported the cued color 98.9% of the time. Participants were instructed not to use any pattern or strategy in choosing between one of the two colors and were told to make a spontaneous decision when the ‘choose’ prompt appeared.

**Figure 1.**
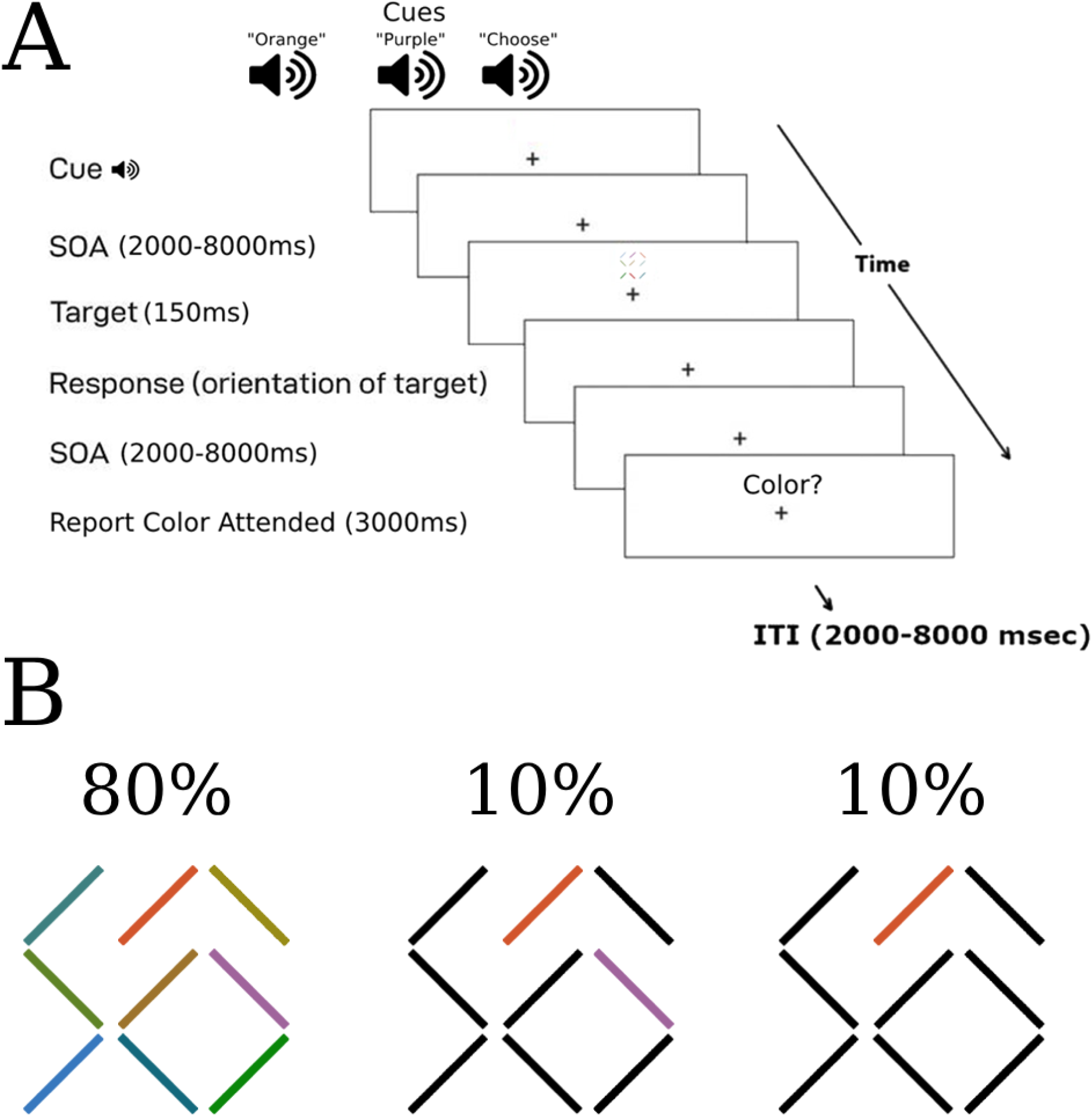
Color willed attention paradigm. **(A)** The cueing paradigm modelled after Bengson et al., 2014, and adapted for willed color attention. Each trial began with an auditory cue (‘orange’, ‘purple’) or prompt (‘choose’), then a variable SOA, followed by the target stimulus (an array of slanted lines of varying colors). Following a variable a second SOA, a report screen appeared displaying the word ‘Color?’ above the fixation point. A variable 2-8 second ITI preceded appearance of the next cue or prompt. **(B)** Example arrays of colored lines for the full nine color arrays (80%), the target color only arrays (10%), and the single color target arrays (10%). The locations and tilt of the various colored lines varied randomly on each trial. The subjects’ task was to discriminate and report the tilt angle of the cued or willed colored line only—except when a single colored line appeared by itself among black lines, in which case they reported the tilt angle of that colored line regardless of whether it was cued or willed.

On 80% of the trials, there was a full array of nine different colored lines, always featuring one orange and one purple line. The orange and purple lines were isoluminant to one another, as verified by a photometer. All colors were developed in DKL space. All stimuli were presented on a background gray screen (RGB: 128, 128, 128). Each line was 0.75 degrees of visual angle in length and 0.75 degrees in width. The whole array of the nine lines subtended 2.75 x 2.75 degrees of visual angle on the monitor. On 10% of the trials, only the two targets (orange and purple) appeared, while the other lines were black. The final 10% of trials were split evenly between only orange appearing (5%) and only purple appearing (5%), with all other lines black. In these conditions participants either responded to the cued or willed color if present, or were required to respond to the unattended color if only that color appeared. During the report period, they were instructed to report the color they chose or were cued, and not the uncued color if it appeared alone. That is, participants were required to report what they had prepared to attend to, rather than the stimulus that actually appeared for the response they made. This manipulation allowed for analysis of responses to validly versus invalidly cued/willed targets, providing a behavioral measure of the effects selective color attention on target processing.

### EEG Recording and Analyses

EEG data were collected from 64 scalp electrodes using a SynAmps II amplifier (Compumedics USA, Inc.) and an actiCAP snap active electrode system (Brain Products GmbH). In accordance with the 10-10 system (Jurcak et al., 2007), electrodes were placed at positions: Fp1, Fz, F3, F7, FC5, FC1, C3, T7, TP9, CP5, CP1, Pz, P3, P7, PO9, O1, Oz, O2, PO10, P4, P8, TP10, CP6, CP2, Cz, C4, T8, FC6, FC2, F4, F8, Fp2, AF7, AF3, AFz, F1, F5, FT7, FC3, C1, C5, TP7, CP3, P1, P5, PO7, PO3, POz, PO4, PO8, P6, P2, CPz, CP4, TP8, C6, C2, FC4, FT8, F6, AF8, AF4, F2 and Iz. Signals were recorded with Curry 8 acquisition software (Compumedics USA, Inc.) at a sampling rate of 1000 Hz. Electrodes at TP9 and TP10 were placed directly on the mastoids of the participants and used as an offline average reference. Impedances were kept below 15 kΩ. Eye-tracking was also conducted using an Eyelink 1000 in remote tracking mode, tracking the right eye of each participant. A tracking sticker was placed on the forehead of all participants to enable remote tracking. A 5-point calibration and validation routine was performed at the beginning of the experiment, with follow-up drift correction performed periodically in between experimental blocks. The eye-tracking was used to ensure that participants were adequately fixating on the designated fixation point.

The paradigm was coded in PsychoPy (Peirce et al., 2019) in Python, and all EEG analyses were conducted in MATLAB, while the reaction time analyses were carried out in R. The data was visually examined and EEG artifacts caused by eye-blinks or eye movements were removed with independent component analysis (ICA) in EEGLAB (Delorme & Makeig, 2004). Artifacts remaining in the data were removed with ERPLAB’s moving window peak-to-peak amplitude function (150 μV threshold) in a 100 ms sliding window moving in 50 ms steps (Lopez-Calderon & Luck, 2014). The data was Fourier transformed to the alpha-band (8-12 Hz) in FieldTrip (Oostenveld et al., 2011) using a tapered convolution with a Hanning taper.

We employed a support vector machine (SVM) pipeline using the fitcsvm() function in MATLAB over each time point, as used in previous decoding studies (Bae & Luck, 2018; Noah et al., 2020). A 3-fold cross validation was used, so the data was equalized across bins (randomly sampled to maintain equal bin lengths), then split into three portions, with two-thirds set aside for training and one-third separated for testing. This was iterated 100 times and the decoding accuracies were averaged across iterations and trials. We used a nonparametric cluster-based permutation technique (Bae & Luck, 2019b), testing significance at each time point against chance (50%) using a one-sample t-test (p < 0.05). This technique controls for autocorrelation in the EEG data by shuffling the class labels when building the null distribution for the cluster mass (iterated 100 times) while maintaining the temporal structure of the data (Bae & Luck, 2019a). For plotting purposes, the data was smoothed over five data points. This process was carried out for each participant.

## Results

### Reaction Time Results

To validate that the paradigm worked as intended, we computed the reaction times to the target in the 10% condition where only one target appeared in the array, which allowed us to assess the effect of attention when a target was the cued/willed color compared to when it was not the cued/willed color. We separated the reaction times based on whether the cue and target matched (valid trial; e.g., cued ‘orange’ and only orange appeared in the array) or did not match (invalid trial; e.g., cued ‘purple’, but only orange appeared). This was done for both the cued (‘orange’ and ‘purple’) and willed (‘choose’) conditions (Figure 2). Repeated-measures analysis of variance (ANOVA) revealed a significant performance difference between valid and invalid trials for both instructed and willed color attention. RTs to validly cued color targets were faster than to invalidly cued targets, demonstrating the behavioral benefits and costs of selective color attention. Separate t-tests showed that valid cued and willed RTs did not differ significantly (p = 0.07). Nor were the invalid (unattended) and willed RTs significantly different (p = 0.88).

**Figure 2.**
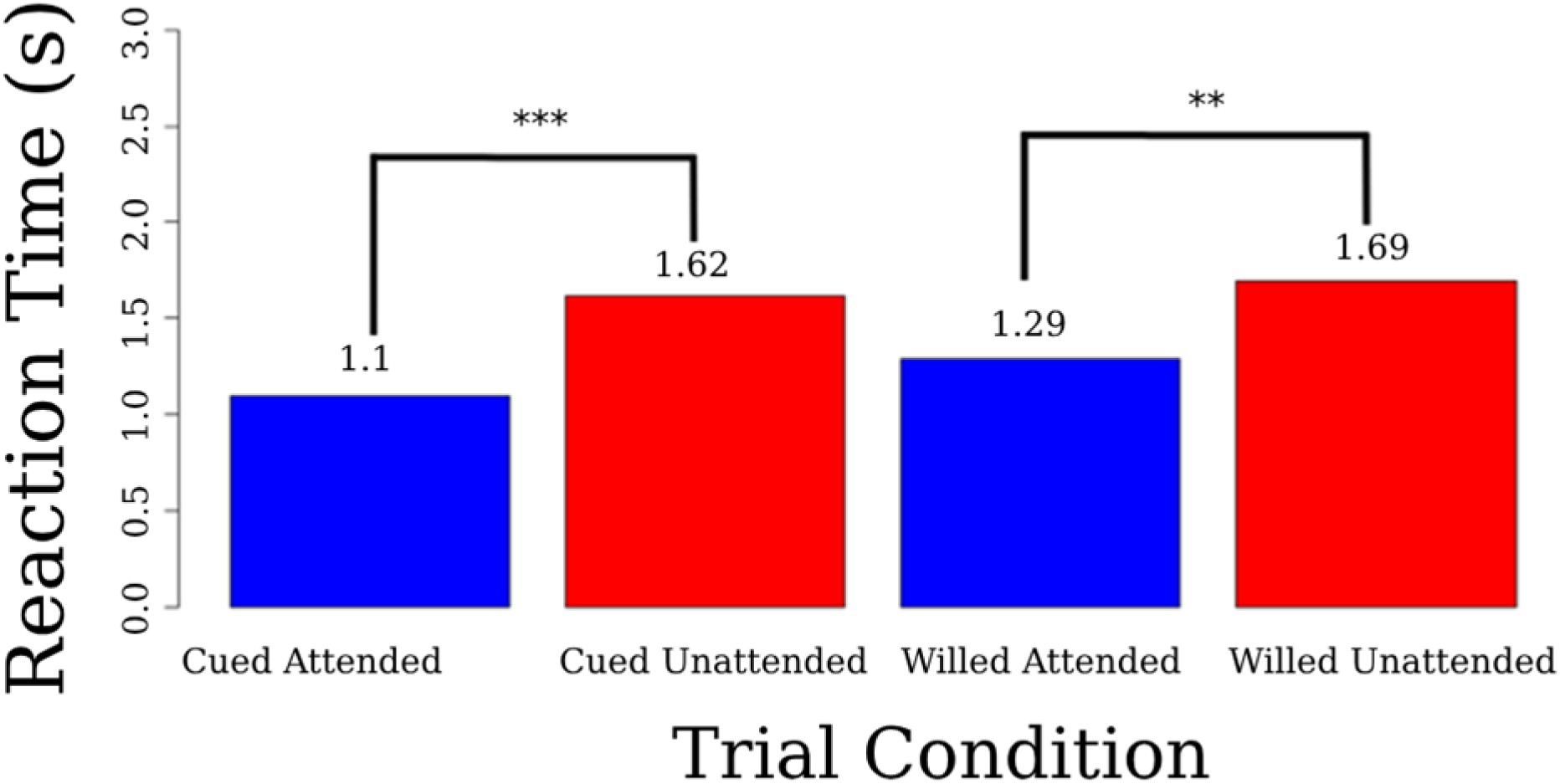
Reaction time validity effects. RTs to the targets in the single color line trials, broken down by cue and willed attention conditions (left side versus right of figure) and cue-target validity (blue bars vs. red bars). ** = p < 0.01, *** = p < 0.001.

To assess whether the individual colors were perceived or treated differently between participants, we also tested RTs to orange versus purple collapsed over all other conditions and found no significant difference (p = .1066). Participants also correctly identified the target’s orientation (left or right) well above chance level (50%) for both orange (94.9%) and purple (91.6%) targets. Finally, to ensure that the selective attention was allocated similarly across the two colors we also tested RT validity effects between orange and purple collapsed across cued and willed attention (Figure 3). Validity effects were significant for both orange and purple colored targets (p < 0.001).

**Figure 3.**
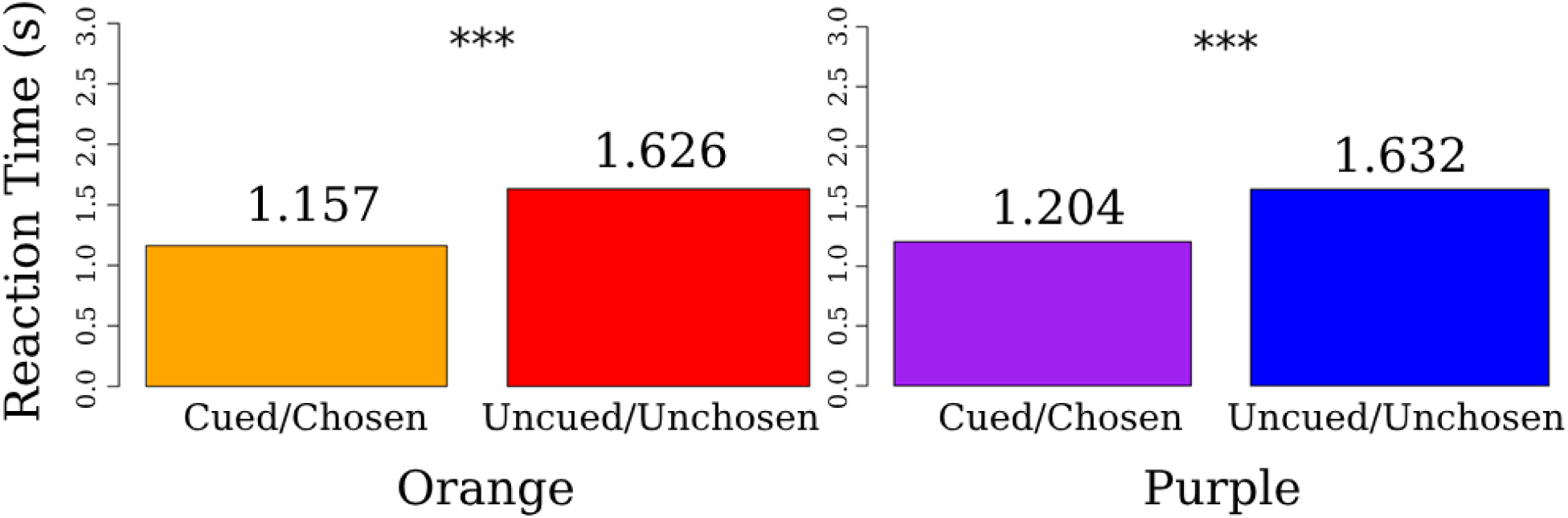
Reaction time effects to targets as a function of color and cue validity in single-line trials. Separate Welch two-sample t-tests were performed to test whether the validity effects were significant for both orange and purple targets. *** = p < 0.001.

### EEG Results

Figure 4 shows the SVM decoding accuracy of the EEG alpha-band power for attend-orange versus attend-purple preceding a choose prompt. Decoding was performed using all 62 scalp electrodes. This result is for the data time-locked to the onset of the auditory cue (verbal word ‘choose’). Decoding in the pre-prompt/pre-decision period is not significantly above chance level. In contrast to the results for willed spatial attention, this pattern suggests that antecedent brain states do not differentiate between color choices (at least with regards to EEG alpha) and therefore do not predict the upcoming choice to be made by the subjects of whether to attend orange or purple.

**Figure 4.**
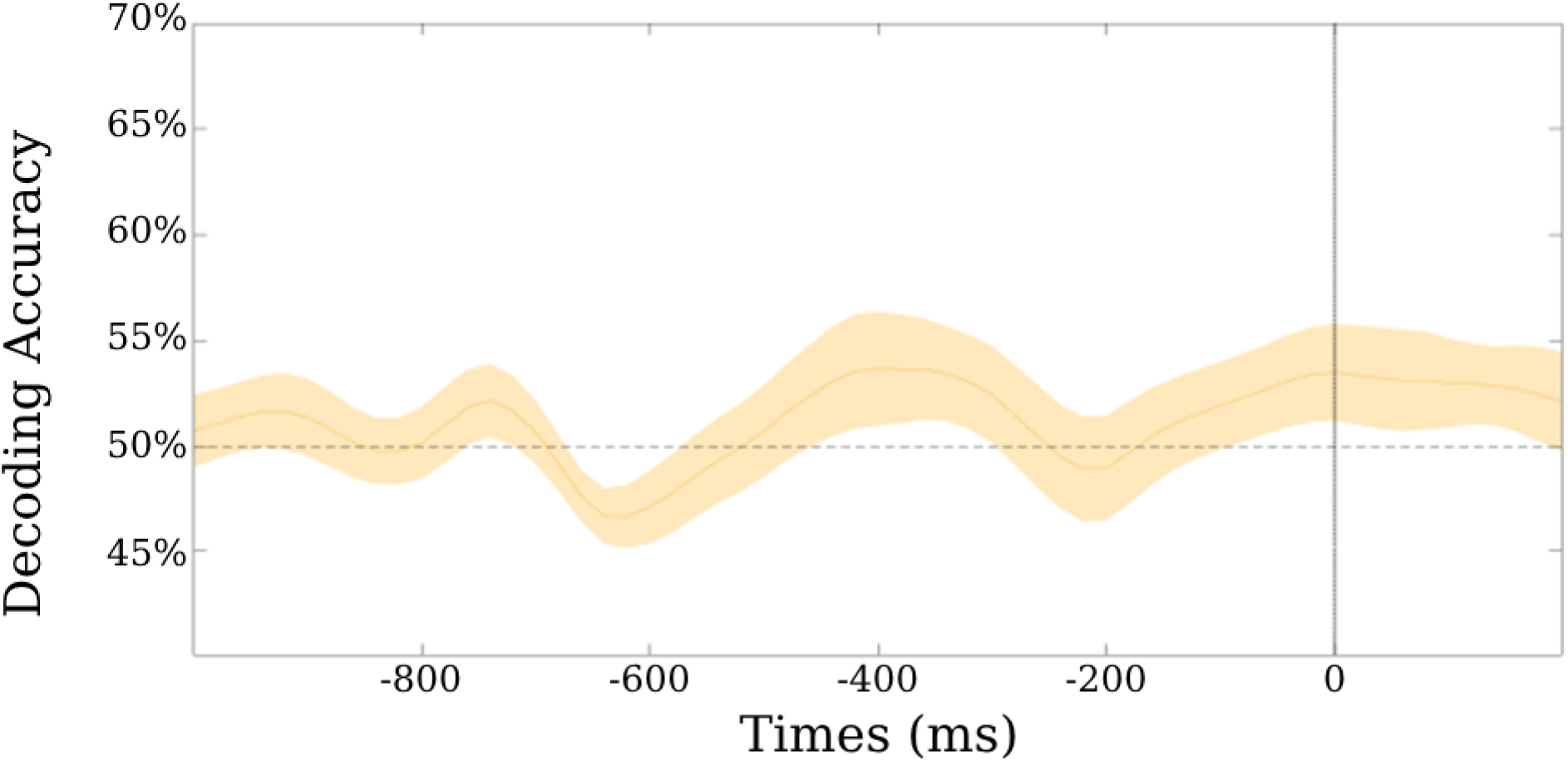
Willed Attention EEG Alpha Decoding in the <u>Pre</u>-Prompt Period. Alpha-band decoding of the pre-prompt period (preceding the ‘choose’ prompt), comparing decisions to attend to orange versus decisions to attend to purple. Unlike prior studies of willed spatial attention, there is no significant decoding in the alpha band prior to the onset of the choose prompt that predicts the subjects’ choices. The black line (t=0 ms) represents the onset of the ‘choose’ prompt.

We also performed EEG alpha decoding over the post-prompt/post-decision period. As shown in Figure 5, significant decoding was obtained after 1000 ms post-prompt onset. Such a pattern could represent differences in the scalp distribution of alpha power as a function of attention to one color versus another, in line with the Gating by Inhibition hypothesis (Jensen & Mazaheri, 2010) and prior results from feature attention (Snyder & Foxe, 2010) and object attention (Noah et al., 2020) studies. The time course of the alpha decoding effects is in line with post-decision occipital alpha lateralization observed in studies of willed spatial attention, which are delayed by approximately 300 ms with respect to those for cued spatial attention, presumably resulting from added decision time in the case of willed attention (Rajan et al., 2018).

**Figure 5.**
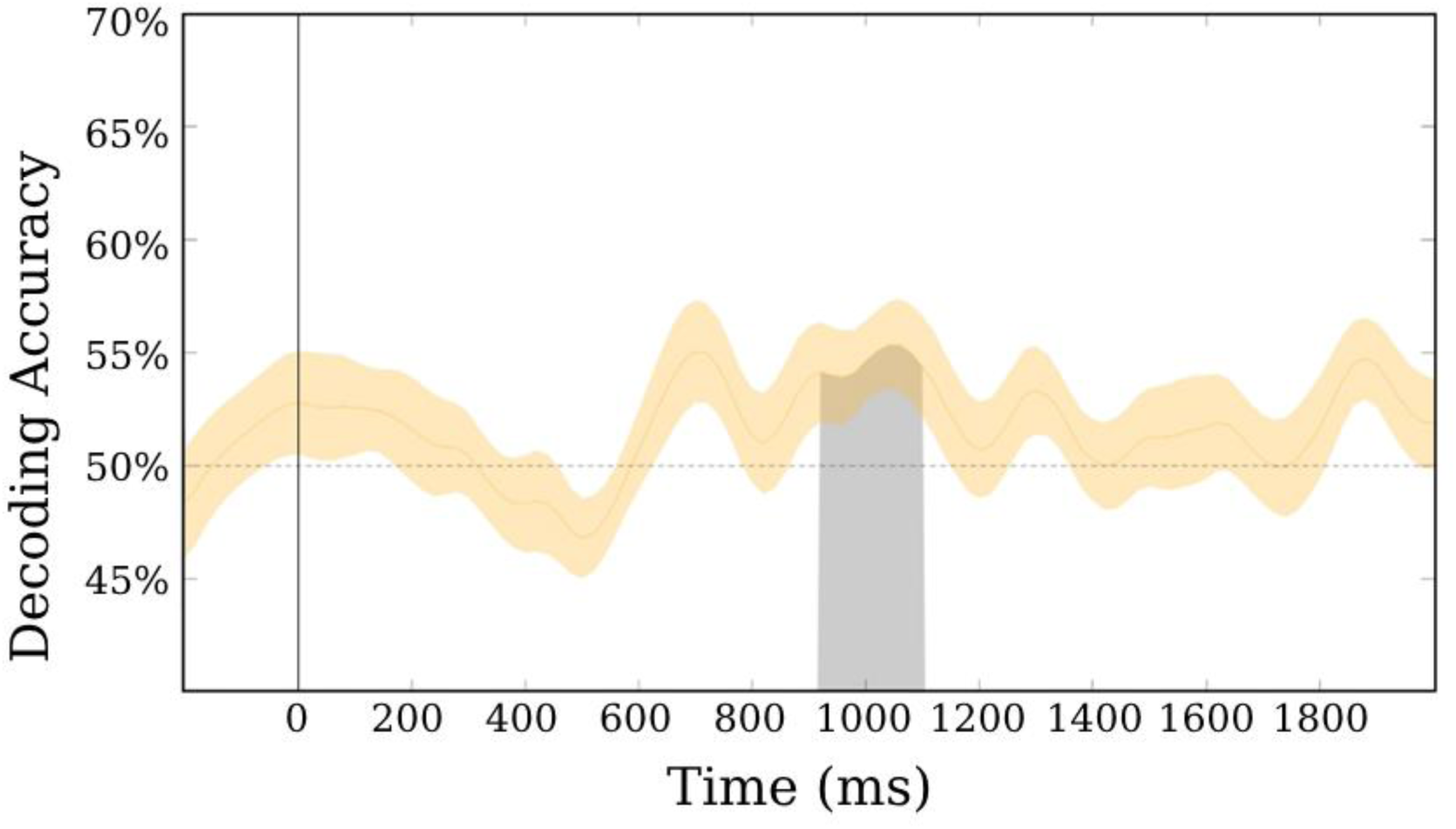
Willed Attention EEG Alpha Decoding in the <u>Post</u>-Prompt/Pre-Target Period. Decoding of the alpha-band oscillations in the prompt-to-target period during willed attention. Comparing decisions to attend to purple versus decisions to attend to orange. Statistically significant decoding was observed between 900-1000 ms after the onset of the prompts. The black line (t=0 ms) represents the onset of the ‘choose’ prompt. Earliest target onset was 2000 ms after prompt onset.

Another possibility is that the alpha difference is not a sign of gating by alpha inhibition differing between colors, but of other binary decisions processes, such as simply choosing one alternative over another. So, rather than differences in alpha gating patterns in visual cortex with attention to different colors, the post-prompt alpha decoding at 1000 ms might reflect differences in task sets, for example (i.e., pick orange vs. pick purple).

To test this hypothesis, we ran the same analysis on the instructed condition, comparing the case where participants were cued to attend to either orange or purple auditorily. If gating by inhibition of one color representation over another explains our alpha decoding, then in line with Rajan et al (2018) we would expect stronger and earlier alpha decoding in the cued attention condition. In contrast to this idea, the analysis showed an even further delay in the onset of significant alpha decoding for being cued to orange versus purple (∼1500 ms after auditory cue onset). This does not seem to support the gating by alpha inhibition view, providing weak negative evidence for the idea that the willed attention effect reflects some sort of binary decision process. Because the cued condition contains twice the number of trials as the willed condition, we can be confident that our failure to observe earlier decoding in the cued condition is not the result of lower signal to noise in the analysis.

We also analyzed the broadband EEG voltage data with the same decoding pipeline to assess whether these findings are strictly alpha-specific. Figure 7 contains the results of the broadband EEG decoding analysis on the ‘instructed’ condition, where decoding began approximately 1,050 ms after the onset of the auditory cue and continued with sporadic patches of significance into 2,000 ms after cue onset. Thus, the electrophysiological correlates of a shift in attention to one of two colors is not limited to the alpha-band (Bengson et al., 2014; Worden et al., 2000). We also conducted the same analysis over the willed condition but found no significant decoding for broadband EEG that predicted subjects’ subsequent choice about what color to attend. Therefore, the post-decision patterns of EEG activity are different for cued and willed attention conditions with respect to EEG alpha versus broadband decoding. The choice process in the willed condition is significantly decodable in the alpha-band but not the EEG broadband, while the inverse is true for the cued color conditions.

## Discussion

Attention in the real world is controlled by competing factors that include voluntary and reflexive attention and their interactions with reward, statistical learning of environmental regularities, and other factors. In this paper we have focused on a pure form of voluntary attention that relies on the free decisions about what to attend, in this case the color of an upcoming visual target. This type of attentional control—willed attention—has primarily been studied in the context of spatial attention (for a review see: Nadra & Mangun, 2023), and willed attention to non-spatial domains has received less attention. Given that color attention mechanisms share some features with spatial attention but also differ in important ways, it is important to investigate the extent to which willed color and spatial attention mechanisms overlap (Andersen et al., 2009; Moore & Egeth, 1998; Saenz et al., 2002; Vierck & Miller, 2007). In this study we used a cueing paradigm to investigate willed color attention in the absence of spatial attention. The task was designed to have high perceptual load, such that using anticipatory attention to prepare for the impending target provided a benefit to the subjects.

We investigated the EEG correlates of attention in time periods prior to and after the arrival of a prompt that indicated to the subjects that they should freely choose what color to attend. In addition, we examined the time period following the instructive cues. In a subset of trials, behavioral measures of color attention were obtained, and showed that the subjects were using the cue or willed information to selectively process the target stimuli. That is, they showed reaction time benefits if the target appeared in the cued or willed color. These behavioral effects of color attention were similar for both cued attention and willed attention.

In the anticipatory period between the cues or prompts and the target, there was significant alpha EEG decoding for the two attention conditions. During the post-prompt period of <u>willed attention</u>, the decisions to attend one of two colors could be decoded in the EEG alpha band (Figure 5). During <u>cued attention</u>, when subjects were directed to attend a specific color, EEG alpha could also be decoded in the cue-to-target period, although at appearing at a slightly longer latency from cue onset (Figure 6). In addition, broadband EEG also showed significant decoding in the post-cue interval (Figure 7), but not in the post-prompt interval during instructed color attention. It is important to note that the willed and instructed conditions had different trial counts because the instructed condition was twice as likely as the willed condition (66.6% instructed/33.3% willed). Thus, the broadband decoding observed following instructive cues but not in the willed color attention condition may be due to higher statistical power in the cued condition decoding analysis.

**Figure 6.**
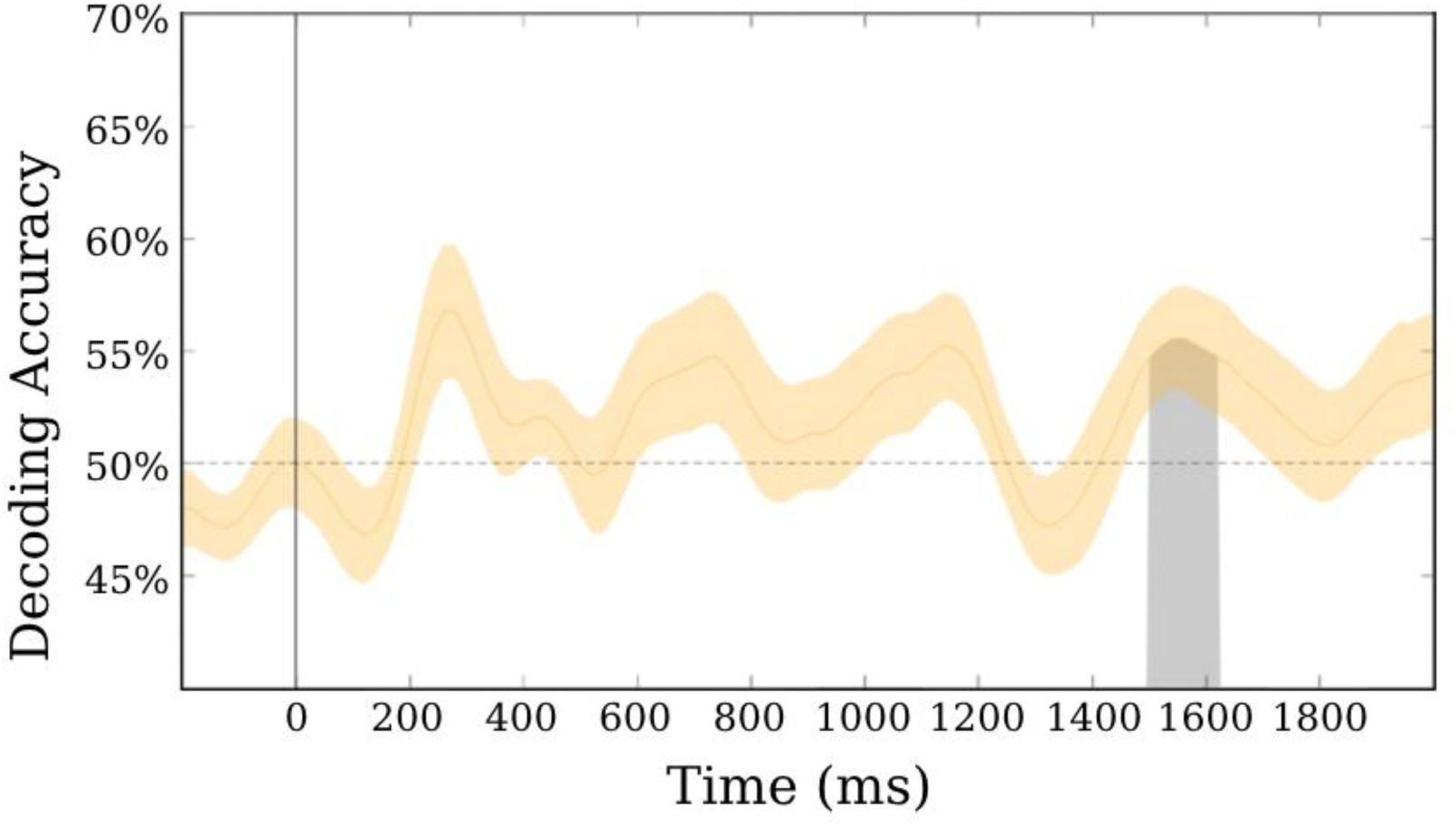
Cued Attention EEG Alpha Decoding in the Post-Prompt Pre-Target Period. Decoding of the alpha-band oscillations in the prompt-to-target period for trials where the subjects were cued (’instructed’) what color to attend (orange versus purple). Statistically significant decoding was observed between about 1500 and 1625 ms after onset of the cues. The black line represents the auditory cue onset. Earliest target onset was 2000 ms after cue onset.

**Figure 7.**
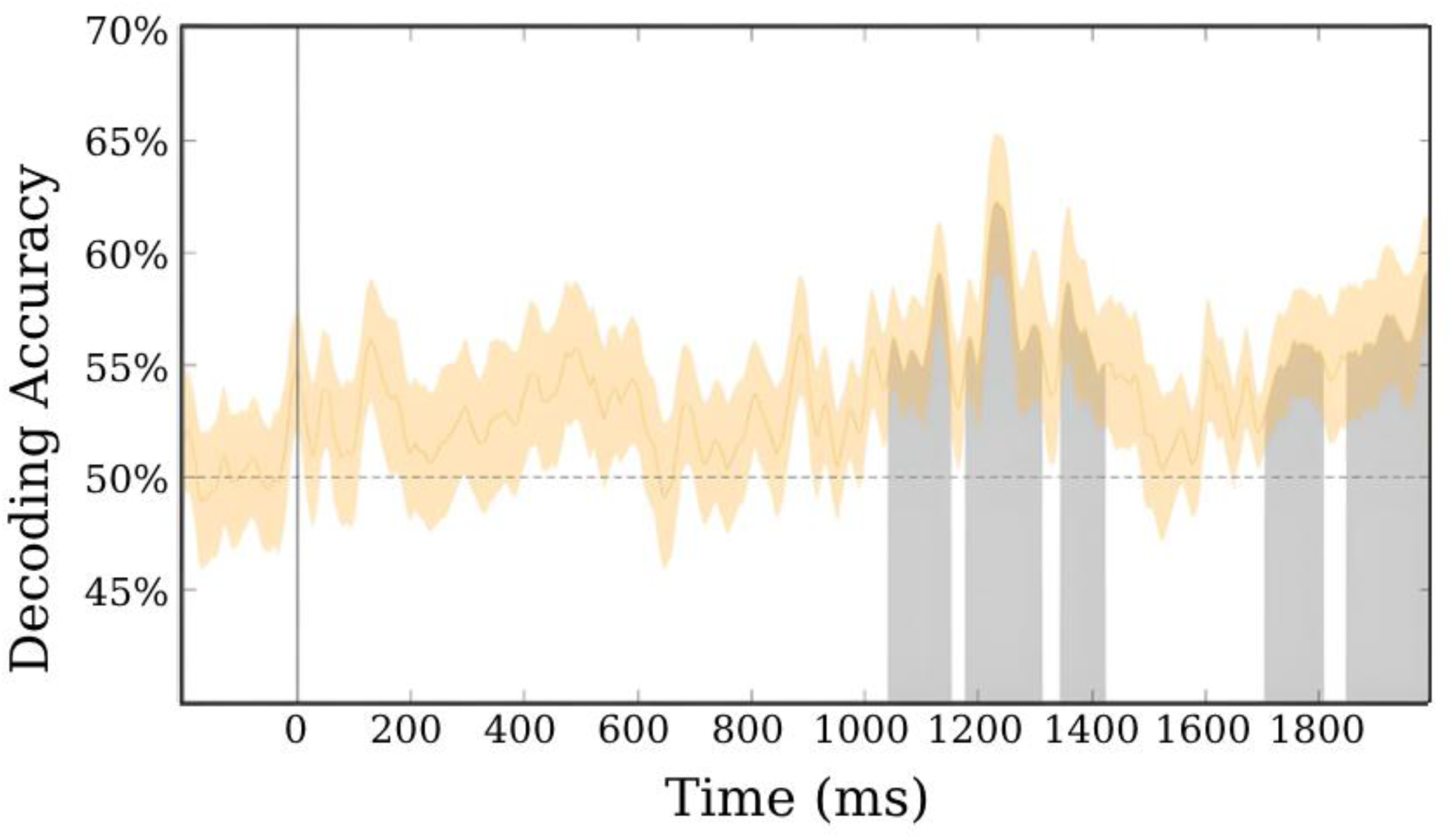
Cued Attention Broadband EEG Decoding in the Post-Prompt Pre-Target Period. Decoding of the EEG broadband activity in the prompt-to-target period for trials where the subjects were cued (’instructed’) what color to attend (orange versus purple). Statistically significant decoding was observed between about 1050 ms to 1400 ms and then again from about 1700 ms to the end of the analysis window. The black line represents the auditory cue onset. Earliest target onset was 2000 ms after cue onset.

We also aimed to test whether the free choices made by the subjects when the prompts appeared could be predicted by the pattern of alpha activity in the pre-prompt period, as is the case for spatial attention (Bengson et al., 2014). Decoding in the EEG alpha band in the hundreds of milliseconds prior to the appearance of the unexpected prompt did not reveal any significant decoding of choices to attend orange versus purple. Thus, we do not find evidence for a shared general purpose mechanism for willed attention. This negative finding replicates Wang and colleagues (2024) using an experimental paradigm that was more similar to that which was used in prior willed spatial attention studies (i.e., Bengson et al., 2014). Together, our findings and theirs converge to suggest that willed spatial attention and willed color attention have different pre-choice patterns of neural activity with respect to the choices made by the subjects. Why might this be?

Prior research on willed attention has focused on spatial attention (Bengson et al., 2015, 2020; Liu et al., 2017; Rajan et al., 2019), specifically on spatial attention between the two visual hemifields. It seems entirely plausible that varying brain states that bias choices about spatial attention between the two visual hemifields, and therefore involving different cerebral hemispheres of the brain, might well be unique to spatial attention. For example, in recent years there has been much evidence showing that the visual field is rhythmically sampled by a shifting spotlight of attention, even when attention is nominally focused on one location (Fiebelkorn et al., 2013; Landau & Fries, 2012; Helfrich et al., 2018). Such a mechanism operating during the pre-prompt period where subjects are simply fixating their eyes on a fixation point and waiting for a cue or prompt might result in a shifting attentional bias to one visual field or the other or time. If such a bias was towards the left visual field in the moment before the prompt appeared, this might result in a choice to attend left on the trial. Additionally, if the bias was towards the right visual field when the prompt appeared, that might result in a choice to attend right. Such a hypothetical mechanism would, therefore, be limited to spatial attention, and would not be expected to generalize to non-spatial willed color attention.

Alternatively, a different mechanism can be envisioned to explain prior willed spatial attention findings for predictive EEG alpha patterns preceding the prompt onset. For example, the signal might not be related to varying lateralized spatial biases shifting attention, but instead could be related to the task structure, which in the spatial attention studies involves the subjects knowing that there are only two spatial locations they are required to choose between on choice trials. These two choices could be represented as a binary decision process (left vs. right) and could reflect a more general binary decision mechanism. If that were the case, this mechanism could just as easily be engaged during willed color attention because in our task (and that of Wang et al., 2024) there were again only two choices (attend orange vs. purple). However, if that hypothesis was correct, pre-prompt predictive EEG alpha signals should have been observed in the present willed color task, but this was not the case.

Another way to consider our findings has to do with the mapping of colors in the visual cortex as compared to the potentially widely spatially separated cortical loci of spatial locations in retinotopic cortex. If the predictive EEG alpha signal observed in prior willed spatial attention studies is related to the Gating of Inhibition Model (Jensen & Mazaheri, 2010), where attended locations show local cortical reductions in alpha power, while ignored locations exhibit local increases in alpha power (representing inhibition), then given the two left and right visual field stimulus locations in the willed spatial attention studies were separated in the brain by a long distance (opposite hemispheres), which would make detection of the differing patterns of EEG alpha on the scalp much easier. In contrast, if the same alpha mechanism was applied to the case of attending to either the orange or purple color, it is likely that the spatial separation in the brain of the differing alpha patterns representing inhibition would be very small. Most work on the localization of color processing in the human brain suggests that such representations are very close together or overlapping in V4 and VO1 (Bartels & Zeki, 2000; Kim et al., 2020; Zeki, 1980; Zeki et al., 1991; Zeki & Marini, 1998).

Therefore, one might not expect the patterns of EEG alpha on the scalp to be different enough to be detected when comparing willed attention to two different colors (orange versus purple). Note, however, we are not advocating that the predictive EEG alpha pattern previously observed for willed spatial attention represents the well-known alpha lateralization that accompanies cued spatial attention because as shown by Bengson et al. (2014, Figure 6), the scalp distributions of the two are somewhat distinct. It is the case, however, that color perception can be decoded from the EEG, and therefore the absence of predictive EEG alpha patterns for willed color attention should not be taken as evidence that EEG is not sensitive to differing patterns of color-related neural activity in the human brain (Chauhan et al., 2023; Hajonides et al., 2021; Laha et al., 2019; Schreiner et al., 2024; Sutterer et al., 2021; Torres-Garcia & Molinas, 2019; Wilson et al., 2023; Wu et al., 2022, 2023).

## Conclusions

In our study of willed color attention, we found no evidence for a pattern of brain activity that predicts what color subjects will choose to attend under free choice conditions. This negative finding is in line with a recent report on color willed attention by Wang et al. (2024). These findings cast doubt on the idea that the predictive EEG alpha patterns previously observed during willed spatial attention represent domain general mechanisms for willed attention. Instead, our findings suggest that willed spatial attention and willed non-spatial (color) attention differ in their neural mechanisms, at least with respect to how antecedent brain states may bias free choices in the context of selective visual attention. Importantly, we observed significant alpha-band power decoding of the willed color after the prompt to decide which color to shift attention occurred. We also found significant alpha-band power and broadband EEG voltage decoding for the instructed condition. These findings indicate that once a decision to shift attention to one of two colors occurs, either volitionally or via external instruction, it is possible to sense the differences in EEG brain signals using support vector machine decoding. Future work should explore how the orienting of non-spatial willed attention is carried out in the brain using functional imaging, which can assess whether similar brain regions are recruited after a prompt to choose in non-spatial attention as was found in prior work on willed spatial attention.

## Notes

### Competing Interest Statement

The authors have declared no competing interest.

